# Counterbalancing the time-dependence effect on the human mitochondrial DNA molecular clock

**DOI:** 10.1101/404582

**Authors:** Vicente M. Cabrera

## Abstract

We propose a new method for estimating the coalescent age of phylogenetically related sequences that takes into account the observed time dependency of molecular rate estimates. Applying this method to human mitochondrial DNA data we have obtained significantly older ages for the main events of human evolution than in previous estimates. These ages are in close agreement with the most recent archaeological and paleontological records.

## Main text

During the last three decades, the application of mitochondrial DNA (mtDNA) variation to studies of human evolution has dominated the scientific scenario. Recently, this small molecule is being substituted by the analysis of whole genomes. However, before turning the page, it would be convenient to solve the patent contradictions between the mtDNA molecular clock time estimations and those proposed by paleontological and archaeological data. A key achievement of early mtDNA analyses was the dating and origin of the most recent common female ancestor of all living women around 200 kya in Africa ^1^. However, in flagrant contradiction, hominin fossils and associated Middle Stone Age artifacts from Jebel Irhoud in Morocco have been recently aged at 315 ± 34 kya ^2,3^. These older dates have been confirmed in a study of ancient African genomes that estimated modern human divergence at 350 to 260 kya ^4^. Another controversial milestone of the mtDNA molecular clock was the dating, based on the coalescence age of macrohaplogroup L3, of the dispersal of modern humans out of Africa 50 to 70 kya ^5^. This is against the presence of early modern human remains in the Levant at the Skhul and Qafzeh caves dated at about 80-120 kya ^6^, the presence of middle stone age industries in the Arabian peninsula with similar dates around 80-130 kya ^7–9^, the recent discovery in southern China of unequivocally modern human teeth dated to 80-120 kya ^10^, or the recently reported detection of an ancient gene flow from early modern humans into the ancestors of eastern Neanderthals more than 100 kya ^11^. Furthermore, the most recent finding of a Homo sapiens maxilla found at Misliya Cave, Israel, dated to 177-194 kya ^12^, could significantly anticipate the exit of homo sapiens from Africa. Curiously, these dates within the MIS-7 last interstadial, are in agreement with the age proposed for an ancient African hominin introgression into the European Neanderthals ^13^. In addition, OSL dating of stratigraphic undisturbed basal stone tool assemblages in Madjedbebe ^14^ placed the human colonization of Australia around 65 kya with minor age uncertainties of only ± 3-4 kyr. The above date is significantly older than the 43-47 kya coalescence age estimated recently from Australian aboriginal mitogenomes ^15^.

The efficiency of the molecular clock ^16^ is based on the reliability of several implicit assumptions as a) the correctness of the mutation rate (μ) of the gene under study; b) constancy with time of the rate of molecular evolution (L) and c) rate homogeneity among the different lineages involved in the phylogeny. To accomplish the first point, in this study we have used the full-length mtDNA germ-line mutation rate of 1.3 × 10^−8^ mutations per site per year (assuming a generation time of 20 years) and its derived rate scalar of one mutation every 4651 years estimated by others ^17^. This mtDNA mutation rate is lower than estimates in most pedigree studies which the authors explain because, at analyzing two tissues, they could discard somatic heteroplasmies. In this respect, it has to be mentioned that second-generation massive sequencing has made possible the direct calculation of the germ-line human genomic mutation rate which resulted in half of the phylogenetic mutation rate. Thus, doubling the estimated divergence dates of Africans and proposing that crucial events in human evolution have occurred earlier than suggested previously ^18^.

It is well known that rates of molecular evolution are not constant neither at interspecific nor intraspecific levels ^19^. In general, they decline with increasing divergence time, but the rate of decay differs among taxa. This time-dependent pattern has also been observed for human mtDNA in coding as well as non-coding regions ^20,21^. Purifying selection on deleterious mutations and mutation saturation have been suggested as the main forces responsible for this time rate decay ^20^. However, the unrealistic large effective population sizes required to explain the long-term persistence of significantly deleterious mutations cast doubts on whether purifying selection alone can explain the observed rate acceleration ^22^. It has also been found unlikely that the apparent decline in rates over time is due to mutational saturation ^20^. Congruently, correcting for the effects of purifying selection and saturation has only slightly modified the mtDNA evolutionary mutation rate, providing molecular times still in apparent contradiction with archaeological and paleontological ages ^20,23^. Demographic processes such as serial bottlenecks and expansions have also been proposed to explain the differences in rate estimation over time ^21^. It seems evident that some adjustment should be implemented to correct the time dependency of the molecular clock. In this paper, we propose a practical approach to counteract the time dependency effect on the molecular rate estimates.

Since early molecular analyses, it was observed that rates of homologous nuclear DNA sequence evolution differ between taxonomic groups ^24^ which was also extensible to mtDNA ^25^. Later, significant differences in rates of molecular evolution between mtDNA human lineages were also detected at haplogroup level ^26–30^. Different relaxed molecular-clock methods have been implemented to incorporate rate variation among lineages ^31,32^. However, the application of these methods to the human mtDNA has yield age estimates for the main milestones of human evolution in agreement with previous molecular estimates^33,34^. In this paper, when distributing mutations of lineages with significant rate differences within coalescent periods, we have used a simple proportionality criterion. We allowed a window from 0 to 5 mutations between sequences as it has been demonstrated that, under a Poisson distribution, even in an extended period of 10,000 years, there could be lineages still carrying the same mutations that their common ancestor, and lineages that have accumulated five new mutations with probabilities higher than 0.05 percent ^35^.

In a population, the fixation time of a mutation (forward) or the coalescence time to the most recent common ancestor (backward), is usually calculated by the estimator 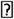 = 4N_e_μ (being N_e_ the effective population size). For the haploid mtDNA genome, 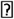 equals to N_ef_μ (being N_ef_ the female effective population size). Although under strict neutral theory parameters, the rate of substitution is equated to the mutation rate ^36^, the same author warned us that a clear distinction exists between mutation rate (μ) and mutation substitution (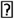). The former refers to the change of genetic material at the individual level, and the latter refers to that at the population level ^37^. Thus, only when N_e_ is constant across generations, keeping small or large sizes, 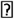 equals to μ. This holds because with large constant sizes the number of new mutations incorporated into the population (Neμ) increases but, on the same path, the probability of fixation (1/Ne) decreases. Contrarily, with small constant sizes, the number of new mutations decreases but the probability of fixation of any of them increases at a similar level. It is widely admitted that N fluctuated largely during the human history and that global exponential population growth is happening since recent times. In this paper, we have taken into account the changes in population size that occurred backward in time and its influence on the rate of gene substitution. When the population size fluctuates across generations, the probability of fixation of a mtDNA neutral variant (q) is no longer the 1/N constant. It will depend on the difference in population size of the next generation (N_1_) with respect to the initial size (N_0_):

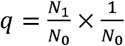

For example, if N_1_ is twice the size of N_0_, q equals 2/N_0_ and, on the contrary, if N_1_ is half the size of N_0_, q equals 1/2N_0_. As a consequence, the rate of substitution for neutral mutations in a population with fluctuating size depends on the change in size between generations:

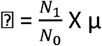

As human populations have been growing exponentially for several centuries, we should counterbalance this effect from the present-day generation (N_n_) going backward in time by inverting the fraction between consecutive generations (N_n-1_/N_n_). Notice that this dependence might explain the differences in rate estimation over time observed empirically (Henn et al. 2009). We will take into consideration this important relationship for the calculation of 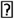.

Several statistics, based on DNA polymorphism, exist to estimate the parameter 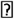. One, S, the number of segregating sites per nucleotide in a sample of sequences ^38^ is strongly dependent on the sample size. A second, π, is defined as the average number of nucleotide differences per site in a sample of sequences ^39^. These two estimators were used to implement a statistical method for testing the neutral mutation hypothesis ^40^. A third statistic, rho (ρ), is referred to as the mean number of nucleotide differences of a sample of sequences compared to their common ancestral type ^41^. This last statistic is calculated from a rooted phylogenetic tree relating the sampled sequences. Although the accuracy of molecular dating with the rho statistic has been questioned because it shows downward biased data estimations, large asymmetric variances and strong dependency of demographic factors ^42^, it is still the most used method to measure intraspecific mtDNA divergence events in humans. In this paper, we propose the use of a modified rho (ρ_m_) that significantly improves the molecular date estimation of key events in the human history based on mtDNA genome data. In order to make explicit our modifications to the classical rho, we have depicted, in figure 1, a real genealogy constructed from five lineages (a), and an idealized star-like phylogeny of the same five lineages (b) supposing a population exponential growth short after a severe bottleneck ^43^. The number of lineages sampled is represented by n_i_; t_i_ are the time periods defined by progressive coalescent events from the tips to the MRCA root; i represents the number of independent lineages left after successive coalescences, 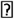_i_ is the number of mutations accumulated during each coalescent period, and m_i_ the number of mutations accumulated along each lineage. Mutations in the star-like phylogeny are distributed into periods following the pattern found in the real phylogeny. As the accumulation of mutations along each lineage is an individual process driven by the mutation rate, μ, and distributed as independent Poisson processes, ρ, the average number of mutations per lineage, has the same value irrespective of the topology. However, as a consequence of the fact that the lineages in the real tree are not independent, because of their shared genealogy, the variance decay is much slower (1/logn) than in the independent star-like tree (1/n) ^44^. As a consequence of this, for the rho calculation, mutations within lineages in the star-like phylogeny are counted only once. On the contrary, in the real phylogeny, only mutations occurring at the tips are counted only once while mutations in subsequent coalescent periods going backward to the MRCA node are counted as many times as the number of the period to which they belong. For this reason, mutations in the older periods are overrepresented in the rho calculation. Because of lineage independence, star-like phylogenies are statistically optimal to calculate ρ and p estimators. Under this topology, as mutations along lineages are counted from the root to the tips in ρ and from tip to tip in π pairwise comparisons, the value of π doubles that of ρ. Thus, to correct for dependence in the real tree, we propose a modified rho statistic (ρ_m_) that is the summation of classical rho statistics calculated for each coalescent period in the tree:

**Figure 1 legend:**
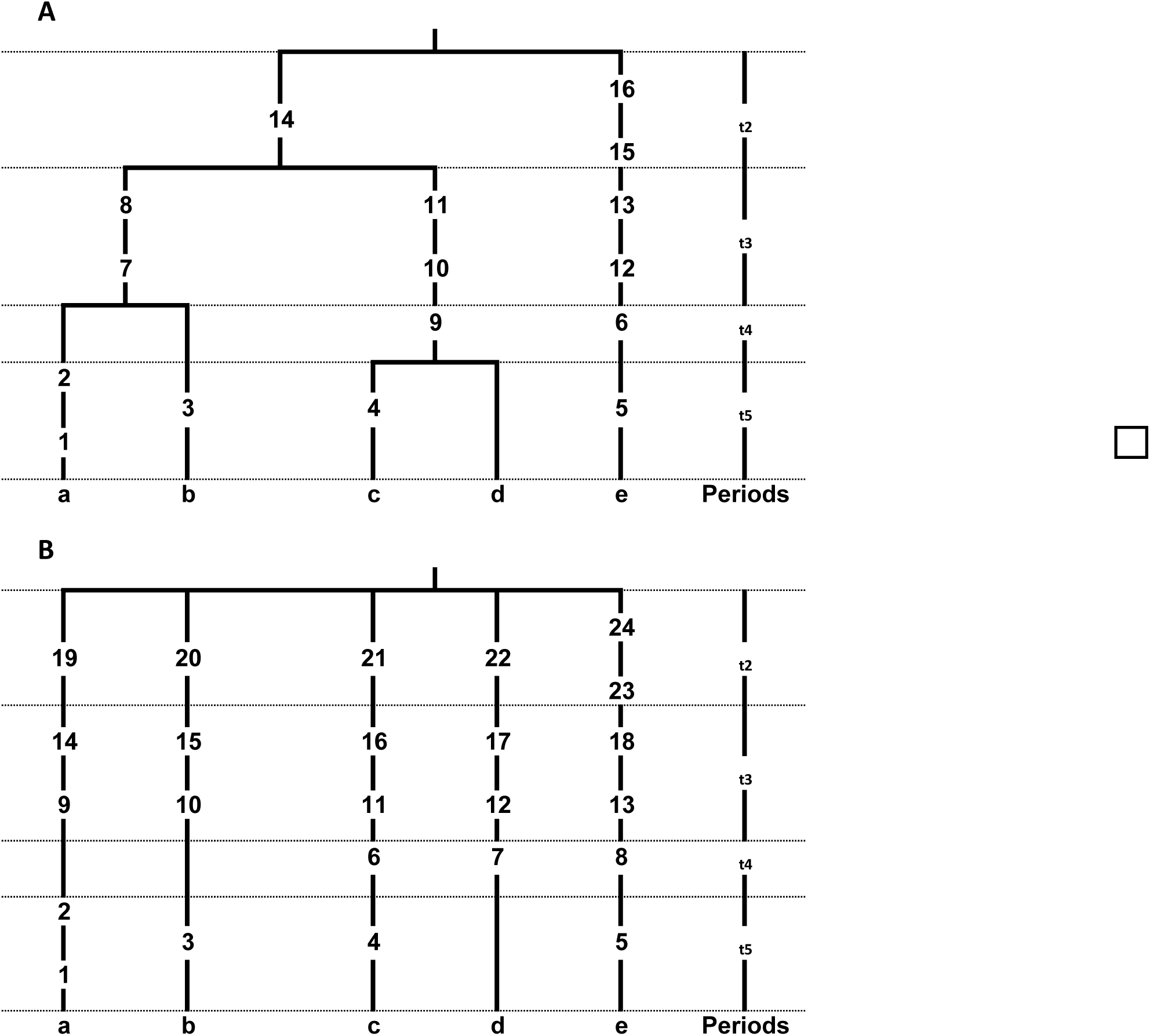
A) Empirical coalescent tree of five lineages (a to e), with four coalescent periods (t) and mutations along branches (numbers). B) Ideal star-like tree of the same five lineages.

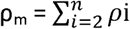

This compound Poisson distribution is also Poisson distributed, therefore mean and variance are equal and, the standard deviation is the square root of this variance. Another important difference between the real and star-like genealogies is that in the first the number of lineages decreases as one stepwise function across coalescent periods while in the second the number of lineages is constant until the root is reached. Equating the number of lineages in the sample to the effective population size in the population, we should take into account this backward real decrease in N_e_ to improve the estimation of the MRCA age. In an ideal coalescent model we should have i-1 population decreasing sizes, but in real phylogenies, in addition to bifurcations, there are also multifurcations and lineages with long internal segments without any branching event. Even so, we applied to each rho in consecutive periods going backward the reverse proportion used to counteract the time dependence effect on the evolutive rate. That is, multiplying in each i-1 period the mutation rate μ by (i-1)/i and leaving the μ rate as calculated from germ-line estimations for the most recent period, comprising the tips of all the lineages sampled. With this method, we have obtained a time-dependent scaled mutation rate, L, that gave human mtDNA intraspecific ages congruent with the archaeological and paleontological calibrated nodes representing key events in the human history.

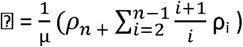

Performing calculations on the empirical genealogy (Figure 1a), we have obtained an age of 22,276.8 ± 149 years using the standard ρ and age of 30,396 ± 174 years, 1.36 times greater, when using the time-dependent L estimator proposed here (Table 1).

**Table 1:** Coalescence age for Figure 1 Tree using the compound rho.

| Period | Lineages | Mutations | Rho | $i/i + 1$ | $\mu$ | $1/\mu$ | Years |
| --- | --- | --- | --- | --- | --- | --- | --- |
| 2 | 2 | 3 | 1.50 | 0.67 | $1.44 \times 10^{-4}$ | 6944 | 10416 |
| 3 | 3 | 6 | 2.00 | 0.75 | $1.61 \times 10^{-4}$ | 6211 | 12422 |
| 4 | 4 | 2 | 0.50 | 0.80 | $1.72 \times 10^{-4}$ | 5814 | 2907 |
| 5 | 5 | 5 | 1.00 | 1.00 | $2.15 \times 10^{-4}$ | 4651 | 4651 |

Coalescence age for Figure 1 tree: $30,396 \pm 5,513$

To apply this statistic to real mtDNA human data we need a rooted tree relating the sampled sequences. Using coalescent methodology we could obtain a probabilistic tree. However, in the case of human mtDNA, we have a very contrasted phylogenetic tree ^45^, constructed using the Network program ^46^. In it, mutations are placed hierarchically from the tips to the root and multiple hits, identified by network reticulations, have been resolved attending to the relative mutation rate of the positions involved ^23^. Thus, following this standard, we constructed an African mtDNA genome-based phylogenetic tree, using 86 previously published complete mtDNA sequences, in which all the main African haplogroups are represented (Figure S1). Likewise, using 142 already published complete mtDNA sequences, we constructed a second phylogenetic tree for the Australasian specific haplogroup P (Figure S2). Finally, to test a more recent human colonization, we constructed a third tree including 48 already published complete mtDNA sequences belonging to the Americas specific haplogroup B2 (Figure S3). Using these trees, we applied our proposed time-dependent based estimator to calculate coalescence ages of several essential nodes of the human history (Tables S1 to S11). We found a TMRCA for all the extant human African mtDNAs of 315,801 ± 17,827 years (Table 2 and S2) which is highly compatible with the recent archaeological and paleontological estimations of modern human origin around 315,000 ya ^2,3^. It has been proposed an early out-of-Africa of modern humans recently, carrying haplogroup L3 precursor lineages, in a favorable time window around 125,000 ya ^47^, that is in harmony with the age calculated here for the L3’4 split in Africa of 165,610 ± 12,869 ya as a lower bound (Tables 2 and S3). Furthermore, this age frame is compatible with the presence of modern humans in the Levant ^6^ and in China ^10^ around 100,000 ya. In the same paper, a return to Africa of basal L3 lineages over 75kya was also suggested. Again, the age for the L3 African expansion calculated with the method reported here of 112,829 ± 10,622 ya makes this suggestion feasible (Tables 2 and S4). Furthermore, under this new temporal window the great morphological variability of the Skhul/Qazfeh remains and their corresponding wide range of ages (120-80 kya), could easily fit into the whole molecular period proposed elsewhere ^47^, beginning with the out-of-Africa of early modern humans and finishing with their early return to the same Continent, carrying basic L3 lineages (125-75 kya). However, notice that a return to Africa from the Arabian peninsula would also be supported by the dates estimated from the archaeological record of the region. On the other hand, our TMRCA for the Austrasia haplogroup P (103,267 ± 10,332 ya) is also in agreement with an early presence of modern humans in Asia (Tables 2 and S6, S7, S8, S9). Thus, it represents a lower bound for the colonization of Philippines ^48^, Sumatra ^49^ and Australia ^14^ around 65,000 to 73,000 ya. It has to be mentioned that from the genome sequencing of an Aboriginal Australian ^50^, it was deduced that Aboriginal Australians are descendants of human dispersal into Eastern Asia that occurred about 62-75 kya. Finally, the age of human expansion into the American Continent, deduced from the haplogroup B2 phylogeny, is around 37,000 ya (Table 2, and table S11). This age is in support of a pre-Clovis occupation of the New World, well before the last glacial maximum. As the calculated ages have wide statistical confidence intervals (Table 2), different models could be adjusted into their frames. For example, we might suppose an earlier out of Africa matching the Misliya maxilla dated to 177-194 kya, then, the Skhul and Qafzeh remains, dated around 80-130 kya might signal the return to Africa of the carriers of the mtDNA basal haplogroup L3 lineages instead of the out of Africa of early anatomically modern humans as proposed here. In the same way, the controversial presence of H. sapiens in Java ^51^ and Sulawesi ^52^ as early as 120 kya would fit into the window age of the Australasian mtDNA haplogroup P (Table 2). Sure, future archaeological discoveries and more precise fossil dating will outline the most appropriate model.

**Table 2:** Coalescence age estimates for several human mtDNA evolutive events.

| Haplogroup split | Evolutive event | Mean age in years | 95% Coefficient interval |
| --- | --- | --- | --- |
| L0/L1'2'5'6'4'3 | Most recent African common ancestor | 317,814 ya | (352,755-282,873 ya) |
| L3'4 | Out of Africa | 165,610 ya | (190,833-140,387 ya) |
| L3 | Return to Africa of L3 | 112,829 ya | (133,648-92,010 ya) |
| P | Reaching the Pacific | 106,752 ya | (127,003-86,501 ya) |
| P | Reaching Australia | 108,034 ya | (128,406-87,662 ya) |
| P | Reaching Philippines | 111,545 ya | (132,244-90,846 ya) |
| P | Reaching New Guinea | 112,070 ya | (132,818-91,322 ya) |
| B2 | Expansion Americas | 37,701 ya | (49,735-25,667 ya) |

